# The impact of overall light-level on the reverse Pulfrich effect

**DOI:** 10.1101/2023.09.27.559782

**Authors:** Victor Rodriguez-Lopez, Benjamin Chin, Johannes Burge

## Abstract

The Pulfrich Effect is an illusion characterized by the misperception of the depth and 3D direction of moving objects. Interocular luminance differences cause the Classic Pulfrich effect; the darker image is processed more slowly. Interocular blur differences cause the Reverse Pulfrich effect; the blurrier image is processed more quickly. A common correction for presbyopia—monovision—intentionally induces the optical conditions that cause the Reverse Pulfrich Effect. The effect sizes, and the fact that tens of millions of people wear these corrections every day, raise concerns about public safety. However, although the impact of overall light-level (e.g., nighttime vs. daytime) on the Classic Pulfrich effect has been well-characterized, its impact on the Reverse Pulfrich effect is unknown. Here, using a custom binocular 4f tunable lens optical system that allows the decoupling of retinal illuminance and retinal blur, we report how the Classic and Reverse Pulfrich effects scale with overall light-level. Both effects increase logarithmically with decreases in light-level. These results motivate a characterization of how light level interacts with other optical factors (e.g., higher-order aberrations) that are likely to impact the Reverse Pulfrich effect, and hence the perceptual consequences of monovision corrections.

**Commercial disclosure:** V. Rodriguez Lopez: None; B. Chin: None; J. Burge: None.

## INTRODUCTION

Catching a frisbee in the hour before night falls is more difficult than during the day. Often, the frisbee arrives before one has had time to react. This phenomenon is explained, in part, by the fact that visual processing is slower when overall light-level is lower. Visual signals that are processed more slowly leave less time for action planning and motor response. The difficulties for vision further increase when the images in the two eyes differ from one another in certain ways. For example, dramatic misperceptions of depth and the 3D direction of motion occur when the image in one eye is, for example, brighter or blurrier than the image in the other eye^1–5^. These illusions are known, respectively, as the Classic and Reverse Pulfrich effects. Signals from the brighter or blurrier eye are processed more quickly than those from the other eye. Signals from the darker or sharper eye are processed more slowly. For moving objects, the differences in processing speed cause effective neural disparities, resulting in the aforementioned illusions.

Changes in overall light-level are commonplace. Over a day, light-level changes markedly, ranging from 10^9^ cd/m^2^ during the day to 10^−4^ cd/m^2^ at night^6,7^. The main purpose of the present paper is to assess how overall light-level interacts with image differences between the eyes to determine the temporal characteristics of neural processing in the binocular visual system. Specifically, we ask how overall light-level impacts the interocular discrepancy in temporal processing caused by a given luminance or blur difference between the eyes. The answer has implications for basic scientific understanding and for clinical practice in ophthalmology and optometry.

Under what circumstances do the images in the two eyes ever differ in retinal illuminance or blur? When the left and right eyes fixate on the same point in a scene, substantial differences in retinal illumination between the left- and right-eye images might occur when viewing specular objects, or when looking at a scene through a pair of sunglasses with one missing lens. Such viewing situations are relatively rare. Substantial blur differences (e.g., ±1.0 D or more), on the other hand, are comparatively common^8,9^. Anisometropia, a condition characterized by interocular differences in optical power of ±1.0 D or more is thought to occur in up to 30% of some important demographic groups^10,11^. And, monovision corrections, which intentionally induce blur differences between the eyes (e.g., 0.75-2.5 D, most commonly around 1.5 D^12^), are being surgically implanted or delivered with contact lenses as popular alternatives to reading glasses, bifocals, and progressive lenses.

For a given overall light-level and focus error, retinal illuminance and defocus blur both vary with pupil size (Figure 1). Thus, to properly investigate these issues, the ability to control focus error and to either monitor or control pupil size is required. We used a custom 4f tunable lens optical system for these purposes. For each combination of overall light-level and focus error, we measured human performance in three fixed pupil-size conditions (2 mm, 4 mm, 6 mm) and one condition in which the pupil was allowed to vary naturally. Each of the fixed pupil size conditions was enforced by projecting diaphragms of appropriate diameter into the pupil plane of each eye. This aspect of the experimental design allowed us to isolate optical from neural influences on the effects. For the natural pupil size condition, the diaphragm was removed such that the entrance pupil of the entire optical system (4f system + human eye) was determined by the diameter of the human pupil. For both participants, the natural pupil size ranged between 4 and 6 mm for all overall light-levels (see Supplementary Figure S1). The 4f optical system included a tunable lens that enabled changes in optical power without inducing magnification differences (Figure 1D). The tunable lens was under programmatic control, allowing us to randomly interleave trials in which the left-eye image was defocused and the right-eye image was sharply focused, and vice versa.

**Figure 1.**
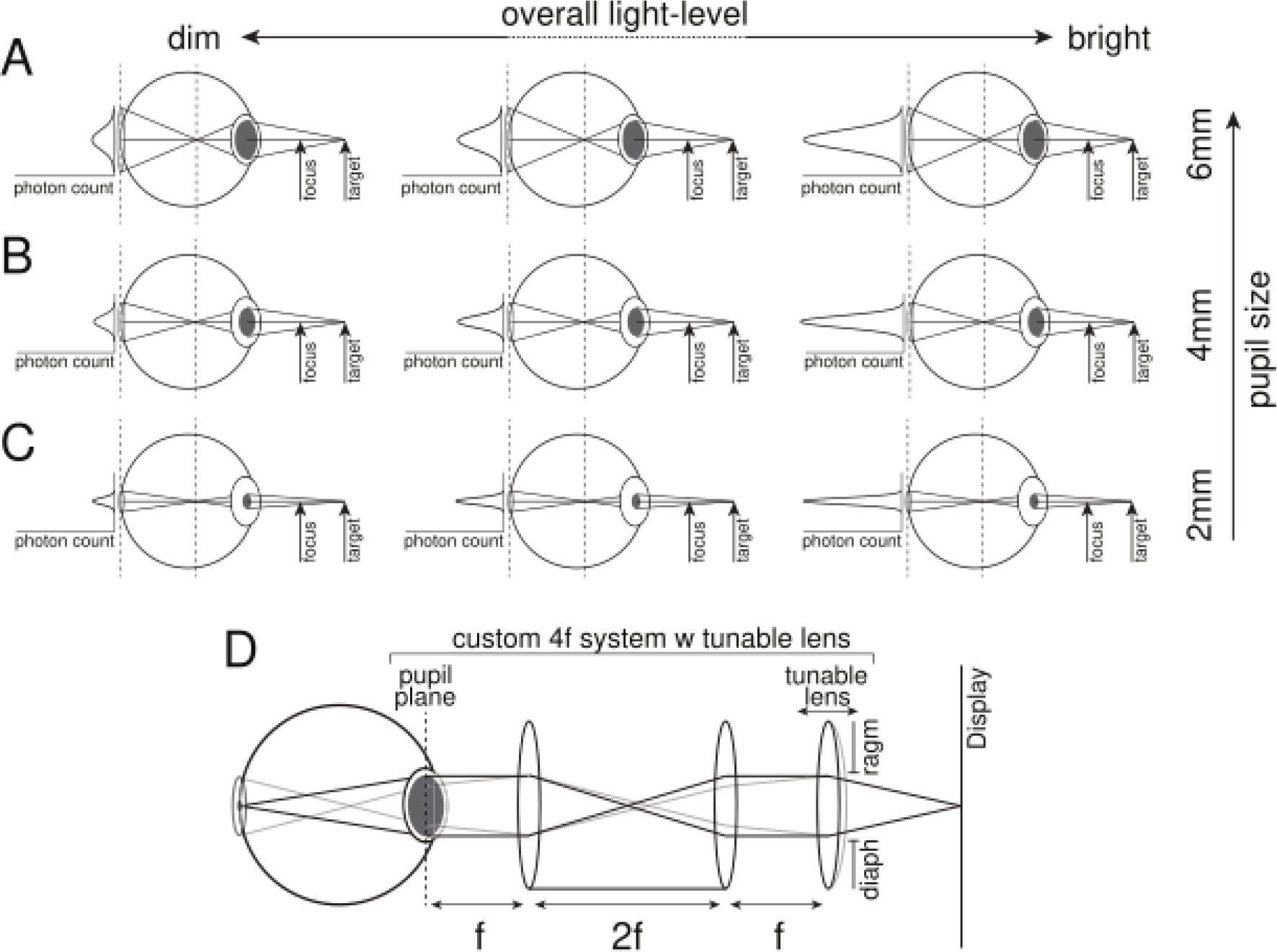
Overall light-level, pupil size, retinal illumination, and optical quality. **A-C**. Effect of changing overall light-level for a fixed pupil size diameter of 6 mm (A), 4mm (B), and 2mm (C). The volume of the photon-count curve represents the amount of light reaching the retina. The width of the curve represents the blur circle. **D**. Custom 4f optical system used in this study. It projects a tunable lens and a diaphragm into the pupil plane of the eye. The power of the tunable lens is under programmatic control.

Here, we investigate how changes in overall light-level change the severity of both the Reverse and Classic Pulfrich effects. With a haploscope system equipped with a 4f tunable lens system for each eye (Figure 2), we measured the effect of changing overall luminance across nearly two orders of magnitude (∼70x), in each of several fixed pupil size conditions and one natural pupil size condition. The effective luminance of the display accounting for light-loss through the 4f system ranged from 12.8 to 0.3 cd/m^2^. Across the luminance and pupil size conditions, retinal illuminance ranged from 360 to

**Figure 2.**
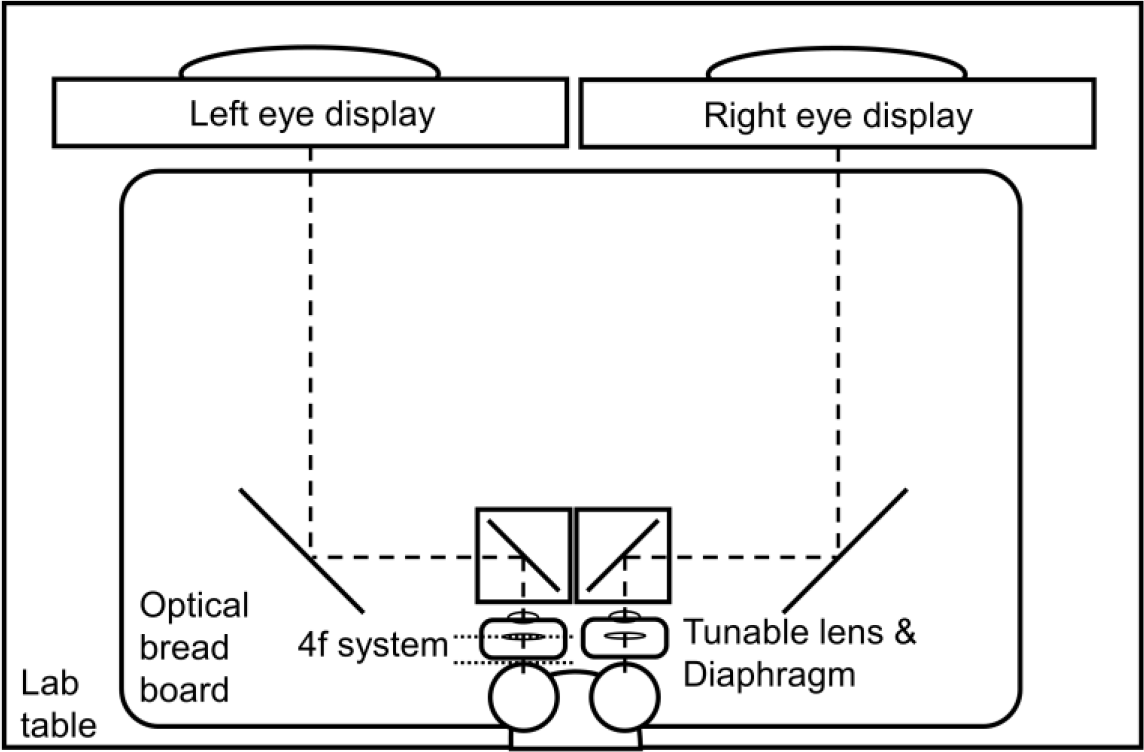
Experimental rig. Representation from above of the optical setup. The 4f optical system shown in Figure 1D is represented as two optical planes. Distances and sizes are not proportional to the real optical system.

0.6 trolands. These are values that straddle the borderline between photopic and mesopic light-levels. The Reverse Pulfrich effect was induced with interocular differences in focus error of 3.0 D^1^, and the Classic Pulfrich effect was induced with interocular luminance differences of 75% (see Methods).

Results show that both the Reverse and the Classic Pulfrich effects increase in strength as overall light-level decreases. The effect of light-level on the Classic Pulfrich effect approximates previous results in the literature^2,13–15^. The effect of overall light-level on the Reverse Pulfrich effect is novel. Results also show that, for matched retinal illuminance, pupil size has a pronounced effect on the strength of the Reverse Pulfrich effect, but it has little effect on the Classic Pulfrich effect. Interestingly, however, the effect of pupil size on the strength of the Reverse Pulfrich effect does not follow simply from geometric optics. We discuss below these findings and their implications. The effect of light-level on the Classic Pulfrich effect is similar to previous results in the literature^2,13–15^.

## METHODS

### Experimental display rig

We used a custom two-display, four-mirror haploscope system, and a prototype portable 4f optical system to present the experimental stimuli (Figure 2). The displays were two identical VPixx VIEWPixx LED displays, with a physical size of 2×29.1 cm, a spatial resolution of 1920×1080 pixels, and a native refresh rate of 120 Hz. To ensure the synchronous presentation of the left- and right-eye images, the displays were daisy-chained together and controlled by the same AMD Radion Pro5300M graphics card with 4GB GDDR6 memory. The light from the displays reached the eyes by first reflecting off a pair of large mirrors and then off a pair of small mirrors. The mirrors were adjusted such that the vergence distance matched the distance of the light path between the monitors and the eyes. The small mirrors were housed in mirror cubes having 2.5 cm diameter circular ports. The mirror cubes were positioned one inter-pupillary distance apart. The effective optical distance of the monitors along the light path from the monitors to the eyes was 80 cm. Thus each monitor pixel subtended 1.58 arcmin of visual angle.

The 4f optical system projected tunable lenses (Optotune, Dietikon, Switzerland) and precision-printed diaphragms of fixed sizes into the pupil planes of the eyes. This system provided a means to programmatically change the interocular focus difference on each trial, and to control the effective pupil size (Figure 1D). A drawback of this system is the substantial loss of light that occurs (∼87%). Through the entire optical system, the maximum luminance was 12.8 cd/m^2^. The optical distance from the eyes of the observer to the screen was 80 cm. Additionally, the baseline power of the tunable lens system was set to +1.25 D to compensate for the distance of the eye to the screen. When the nominal defocus was set to 0.0 D, this baseline power prevented subjects from having to accommodate to clearly focus the display.

As noted, the custom 4f tunable lens systems were portable prototypes. The lens systems were in development for a commercial head-mounted ophthalmic device for simulating multi-focal corrections (SimVis Gekko, 2EyesVision, Madrid, Spain). This system was optimized for head-mounted device development and not for conventional psychophysical experimentation on an optical bench. As a consequence, a calibration procedure was required before data was gathered to ensure that the center of optics of each subject’s eye and each 4f tunable lens system were properly aligned. This procedure required extensive comparisons of subtle visual markers (vignetting, aspect ratio, tilt, dynamic blur, etc.) and entailed that it was impractical to collect data from naïve observers (see Discussion).

### Stimuli

The stimulus consisted of four strips textured with randomly positioned 1.00×0.25 degrees white bars moving horizontally at a constant speed of 4.0 degrees per second. Adjacent strips moved in opposite directions (Figure 2).

To induce an effective onscreen interocular temporal shift Δ*t* (i.e., delay or advance), we first determine the equivalent onscreen spatial disparity in degrees of visual angle

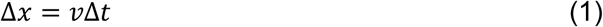

where *v* is the movement speed in degrees per second. The onscreen horizontal left- and right-eye positions of the strips in the two eyes evolve with time according to

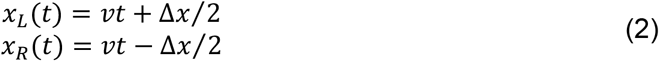

where *x*_*L*_ and *x*_*R*_ are the left- and right-eye x-positions in degrees of visual angle, and *t* is time.

When the onscreen interocular temporal shift equals zero, the strips are specified by onscreen spatial disparity to move in the plane of the screen. When the onscreen interocular temporal shift is non-zero, the strips are specified by onscreen spatial disparity to be in front or behind the screen. Negative interocular temporal shifts indicate that the left-eye onscreen image is delayed relative to the right-eye image. Positive interocular temporal shifts indicate that the left-eye onscreen image is advanced relative to the right-eye image. For negative temporal shifts, rightward and leftward moving strips are specified by disparity to be farther than and nearer than the screen plane, respectively. For positive temporal shifts, rightward and leftward moving strips are specified by disparity to be nearer than and farther than the screen plane, respectively.

### Procedure

The task was to set the effective onscreen temporal shift (i.e., the onscreen spatial disparity) via an adjustment procedure until all strips appeared to move in the plane of the screen (Figure 3). In each condition, six runs were conducted. On a given run, the initial onscreen interocular temporal shifts ranged from -15 ms to +15 ms. On a given trial within a run, the strips moved continuously with a fixed temporal shift until the observer made either a coarse adjustment (±1.0 ms) or a fine adjustment (±0.2 ms). Each adjustment initiated the next trial, with a new onscreen temporal shift. The observer continued running trials and adjusting the onscreen temporal shift until the observer indicated with a button press that the task had been completed. Throughout each trial, the observer fixated on the rightward-moving strip nearest the center of the screen. Sometimes, the rightward moving strip was just above the vertical midpoint of the screen; sometimes it was just below the vertical midpoint. The stimulus texture was updated with a new texture (different random configuration of the bars within the moving strips of the stimulus) in every run.

**Figure 3.**
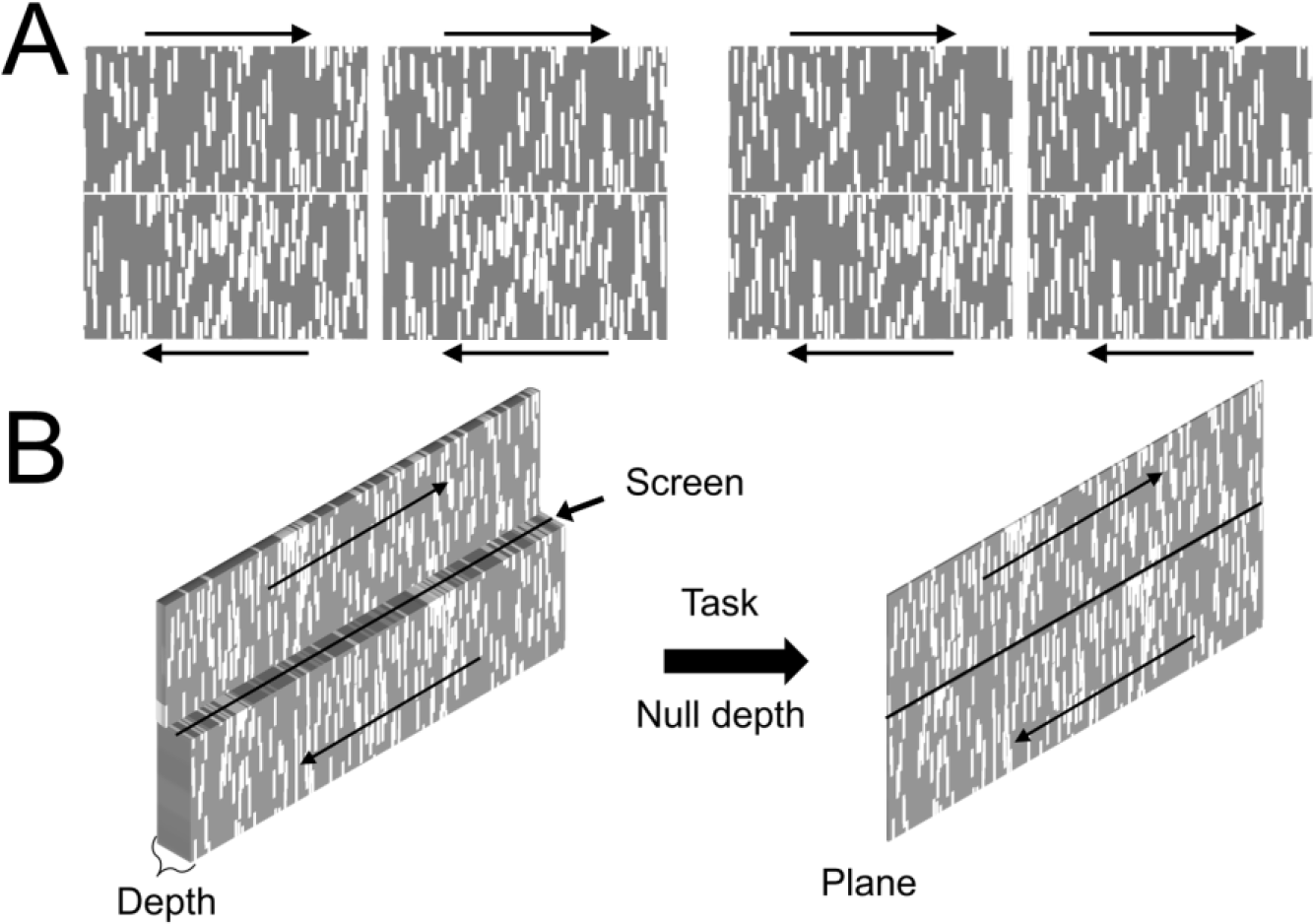
Stimulus and the depth nulling task. Adjacent strips moved in opposite directions. **A**. Head on view. On the left, left- and right-eye half-images of a stimulus with a disparity-specified depth step. Free-fuse to see the disparity-specified depth. Divergent-fusing will make the top strip appear farther than the bottom strip. Cross-fusing will make the top strip appear closer than the bottom strip. On the right, left- and right-eye half-images of a stimulus with zero disparity between the images that specifies a flat plane. **B**. Perspective view. On the left, the upper strip is perceived to be behind the screen, and the bottom strip is perceived in front of the screen. On the right, both the upper and lower strips are perceived to be in the plane of the screen.

The critical onscreen delay for a given condition was estimated by averaging the final settings of the onscreen delay across the six runs. The critical onscreen delay indicates the onscreen delay required to null the neural delay caused by a particular interocular difference in defocus blur or retinal illuminance. When the critical onscreen delay is negative, the left-eye image has to be delayed onscreen relative to the right-eye image for the bars to appear to lie in the same depth plane. When the critical onscreen delay is positive, the left-eye image has to be advanced onscreen relative to the right-eye image. The standard deviation of the final settings indicates the uncertainty with which the final settings were set. It is clear from a cursory examination of the data that the effect sizes are much larger than measurement uncertainty.

### Overall light-level

To compute the overall light-level in the context of the current experiments, the luminance output of the monitors, the light loss along the optical path to the eyes, and the effective pupil size must all be determined. First, we measured the maximum luminance output of the monitors using a spectroradiometer (120 cd/m^2^). Next, we measured the effective luminance of the monitors through the custom 4f system and verified that the measurements agreed with calculations indicating the expected light loss. Light loss due to the custom 4f system was high (∼90%), so the maximum effective monitor luminance was 12.8 cd/m^2^.

Stimuli were presented at four overall luminance levels ranging from 12.8 cd/m^2^ (maximum luminance reaching the eye) to 0.2 cd/m^2^. To reduce the luminance from the maximum level, we positioned a neutral density filter into the light path for each eye. The neutral density filters were always matched between the eyes and had optical densities (OD) with one of four values (0.0, 0.6, 1.3, and 1.8). These optical densities correspond to transmittances of 100%, 25%, 5%, and 1.5%; the three darkest overall light levels correspond to 4x, 20x, and 67x less light than the brightest condition.

### Pupil size

Pupil size was controlled by projecting a diaphragm of known size into the pupil plane of each observer using the custom 4f-optical system referenced above (Figure 2). Cycloplegic drops—2 drops of phenylephrine 2.5% and 2 drops of tropicamide in each eye—were administered to dilate the pupil and ensure that the entrance pupil diameter was determined by the projected diaphragm. Measurements were taken with fixed pupil diameters of 2 mm, 4 mm, and 6 mm. Measurements were also taken with natural pupil diameter. For the natural pupil, no cycloplegic drops were administered and a diaphragm with a 10 mm diameter was inserted into the 4f-optical system such that the entrance pupil of the optical system was determined by the natural pupil of the observer’s eye.

To obtain measurements of the natural pupil size, posthoc measurements were taken under the exact same stimulus and lighting conditions as the main experiment. Observers wore Pupil Core 120 Hz camera glasses (Pupil Labs, Berlin, Germany), a mobile eye-tracking system that also provides pupillometry. Pupil diameter data were obtained for 1 sec (i.e., 120 frames) after 3 seconds of adaptation to the light level. We report the average across frames as the pupil size for a given stimulus condition. The results indicated that the natural pupil ranged in size from approximately 4 mm to approximately 6 mm, from the highest to the lowest overall light-level (see Supplementary Figure S1).

### Experiments

The conditions were grouped into two different experiments. The first measured the Reverse Pulfrich effect. The second measured the Classic Pulfrich effect.

The Reverse Pulfrich effect was measured by inducing focus error in one eye while leaving the other eye unperturbed. The interocular blur difference in diopters is given by the difference between the focus error induced in the right eye minus the focus error in the left eye

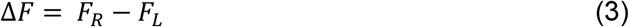

For each overall light-level, we measured two conditions. In one, the left eye was defocus blurred and the right eye was sharp (Δ*F*=-3.0 D). On the other, the right eye was defocus blurred and the left eye was sharp (Δ*F*=+3.0 D). These defocus differences were induced by increasing the power of the tunable lens in front of either the left eye or the right eye, thereby setting the optical distance of the corresponding monitor beyond infinity and causing retinal defocus blur. The retinal defocus blur difference, assuming geometric optics, can be calculated from the diopters of defocus and the pupil size as follows

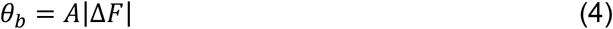

where *θ*_*b*_ is the blur diameter in radians, *A* is the pupil aperture (diameter) in meters, and |Δ*F*| is the interocular difference in defocus.

The Classic Pulfrich effect was measured by reducing the luminance of one eye’s image onscreen while leaving the other eye unperturbed. The interocular luminance difference, expressed in units of optical density (OD) is given by the difference between the effective optical density associated with the right-eye image minus the effective optical density on the left-eye image

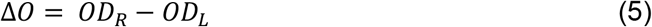

For each overall light-level, we measured two conditions. In one the left-eye image had lower luminance than the right (Δ*O*=-0.6 OD). In the other condition, the right eye image had a lower luminance than the left (Δ*O*=+0.6 OD). These conditions correspond to the left eye receiving 25% of the light that the right eye receives, and vice versa.

The retinal illuminance for each overall luminance level and pupil size was computed by multiplying the luminance by the pupil area expressed in mm^2^

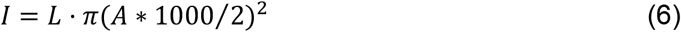

where *l* is the retinal illuminance in trolands, *L* is the luminance level in cd/m^2^, and, again, *A* is the pupil aperture (diameter) in meters.

The estimated delay, and standard error of the estimated delay, at each retinal illuminance level were calculated, respectively, by averaging the final adjustment settings and by taking the standard deviation of the final adjustment settings divided by the square root of the number of settings.

To fit how the onscreen delay changed with retinal illuminance, we used a power function

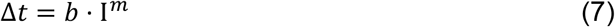

where Δ*t* is the delay, *I* is the retinal illuminance, and *m* and *b* are the parameters of the power law function that are adjusted to minimize the square error of the fit. Taking the natural logarithm of both sides of the equation shows that, in log-space, the power law function has the equation of a line

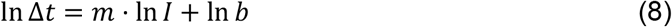

We transformed the data by taking the natural logarithm of both onscreen delays and the retinal illuminances and then fit the logged data with least squares regression, weighted by the standard deviation of the final adjustment settings. Performing the fit in log-space was justified because the standard error of the final interocular delay settings across runs was more nearly constant on a log scale than on a linear scale.

Each human observer ran in a total of 64 conditions (i.e., 4 overall light levels x 4 pupil sizes x 2 retinal illuminance differences x 2 defocus differences) for each experiment. Overall, 16 retinal illuminance levels ranging from between 0.6 to 360 trolands were measured. The range of light-levels varied between photopic and mesopic conditions. The whole experiment took approximately 3 hours to complete.

## RESULTS

We measured the impact of overall light-level on temporal processing in the visual system by taking advantage of both the Reverse and Classic Pulfrich effects.

Two experiments were run. The first experiment assessed how changes in overall light-level changed the strength of the Reverse Pulfrich effect, which is induced by interocular differences in focus error. The second experiment assessed how changes in overall light-level changed the strength of the Classic Pulfrich effect, which is induced by interocular differences in the amount of light entering each eye. In both experiments, pupil size was either fixed to one of three diameters (2 mm, 4 mm, and 6 mm), or it was measured as it varied naturally during the experiments (see Methods). Across the four overall light-levels (i.e., display luminances) used in the experiment, the natural pupil sizes ranged from between 4 mm to 6 mm (see Supplementary Figure S1). In total, there were 16 distinct retinal illuminance levels ranging from between 0.6 to 360 trolands.

Subjects viewed four horizontally drifting strips textured with vertical bars, that were stereoscopically specified to be in front of, in line with, or behind the plane of the screen. The task, in an adjustment procedure, was to adjust the apparent depth until all strips appeared to be moving in the plane of the screen. Neighboring strips always drifted at the same speed in opposite directions (left vs. right), and the onscreen disparity associated with adjacent strips was always equal in magnitude and opposite in sign. The task was intuitive and easy for subjects to perform.

In the Reverse Pulfrich experiment, for each overall light-level, data was collected in each of two conditions. In one condition, the left-eye image was more defocused (Δ*F* = -3.0 D) and hence blurrier than the right-eye image; in the other condition, the right-eye image was more defocused than the left-eye image (Δ*F* = +3.0 D).

Figure 4A shows the onscreen delay for the 12 adjustment runs (6 runs x 2 conditions) from one observer at a particular overall light-level and pupil size (0.2 cd/m^2^ and 4 mm). (This data is representative of the raw data in other conditions.) The average across the final settings of all six runs in a given condition provides an estimate of the critical onscreen delay that was required to make the drifting bars appear to move in the plane of the screen. This critical onscreen delay should be equal in magnitude and opposite in sign to the neural delay caused by the interocular difference in light level. When the left eye was blurry, the critical onscreen delay was -11.4±4.9 ms (ΔF=-3.0 D). When the right eye was blurry, the critical onscreen delay was 11.7±3.6 ms (ΔF=+3.0 D). These results indicate that the image in the manipulated (blurrier) eye was neurally processed more quickly. These results are consistent with previously reported results^3,4^. Thus, blurring an image speeds up how quickly that image is processed by the visual system.

**Figure 4.**
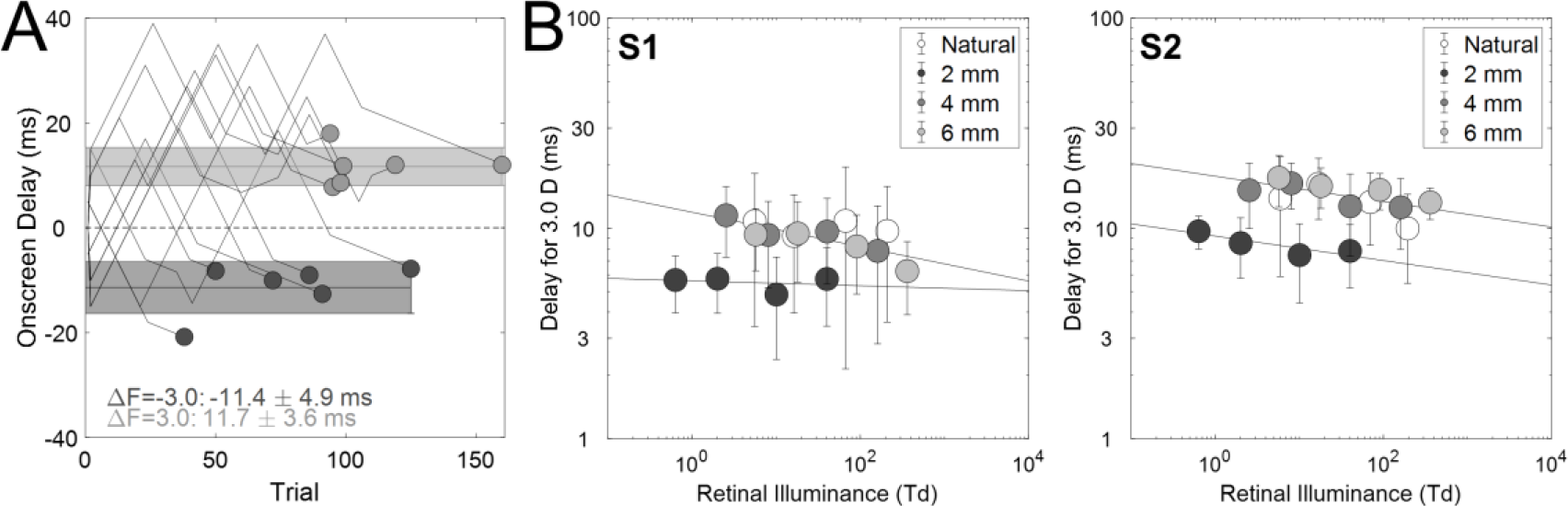
Reverse Pulfrich effect results. **A**. Onscreen delays presented to a subject during each of six adjustment runs in two separate conditions (jagged lines). The task was to adjust the onscreen delay until the strips appeared to move in the plane on the display (dots). Each run had a different random starting point between -15 ms and 15 ms. Negative onscreen delays indicate that the left-eye image was presented later on-screen than the right-eye image. Positive onscreen delays indicate left-eye image was presented earlier on-screen. Runs from the condition in which the left-eye image was blurred by +3.0 D of focus error and the right-eye image is sharply focused are in dark gray (*ΔF* = -3.0 D). Runs from the condition in which the left-eye image was sharply focused, and the right-eye image was blurred by +3.0 D of focus error are in light gray (*ΔF* = +3.0 D). The dashed dark gray and light gray lines represent the average of the final settings in each condition. The shaded dark gray and light gray regions represent the standard deviation of the final settings in each condition. **B**. Critical onscreen delay for all combinations of retinal illuminance level (abscissa) and pupil size (colors) for subject 1 (left subplot) and subject 2 (right subplot). Critical onscreen delay increases when the retinal illuminance decreases, and pupil size has no notable effect. For the same focus error, the smallest pupil size (2 mm) produced notably smaller effects than other pupil sizes. The Reverse Pulfrich effect is stronger when light-level is lower.

Figure 4B shows the impact of overall light-level on the strength of the Reverse Pulfrich effect, one panel for each observer. Each data point is the (average) magnitude of the critical onscreen delays in the two defocus conditions at a given light level and pupil size (see Fig. 4A). Larger interocular delays are associated with lower light levels, smaller interocular delays are associated with higher light levels, and the change in interocular delay with light-level is approximately linear on a log-log scale. For S1 (Figure 4B, left panel), the slope of the linear regression in the log-log space reflects the power of the power function, which is -0.08 ms/td (CI^68^ = [-0.09, -0.04]) for natural, 4 mm and 6 mm pupil sizes and -0.01 ms/td (CI^68^ = [-0.05, 0.004]). For S2 (Figure 4B, right panel), the slopes are -0.06 ms/td (CI^68^ = [-0.09 -0.04]) for natural, 4 mm and 6 mm pupil sizes and -0.06 ms/td (CI^68^ = [-0.09, -0.03]) for 2 mm pupil size.

Note that the interocular delays associated with fixed 2 mm pupils are quite a bit smaller than the interocular delays associated with larger pupil sizes. This is to be expected. A given focus error (e.g., 3.0 D) produces less retinal blur when pupil sizes are small (Equation 5), so the difference in retinal blur between the eyes, and hence the interocular delay, is expected to be smaller. However, by the same reasoning, it is a bit surprising that there are no clear differences among interocular delays associated with the fixed 4 mm pupils, fixed 6 mm pupils, and natural pupil sizes which ranged from 4 mm to 6 mm. We speculate that this pattern in the data can be accounted for by the presence of higher-order aberrations in the human eye. We discuss this possibility in the Discussion section below.

In the Classic Pulfrich experiment, for each overall light-level, there were again two conditions. In one condition, the left-eye image was dimmer and received only 25% of the light that the right eye did (ΔO = -0.6 OD). In the other condition, the right-eye image was dimmer than the left eye (ΔO = +0.6 OD). Six adjustment runs were completed for each condition. (Note that an optical density of 0.6 corresponds to a 25% transmittance, which is equivalent to a 75% light-loss).

Figure 5A shows all 12 adjustment runs (6 runs x 2 conditions) from one observer at another overall light-level and pupil size (3.2 cd/m^2^ and 6 mm). This data is representative of the raw data in other conditions. At this light level, when the left eye was dark (ΔO =-0.6 OD) the critical onscreen delay was +12.6±1.0 ms, indicating that the dark left-eye image had to be advanced onscreen to counteract the fact that it was neurally delayed. When the right eye was dark (ΔO =+0.6 OD), the critical onscreen delay was -12.4±0.9 ms, indicating that the left-eye image had to be delayed onscreen to compensate for the fact that the dark right-eye image was neurally delayed (Figure 5A). Unlike in the previous experiment in which the manipulated (blurrier) image was neurally processed more quickly, in this experiment the image in the manipulated (dimmer) eye was neurally processed more slowly.

**Figure 5.**
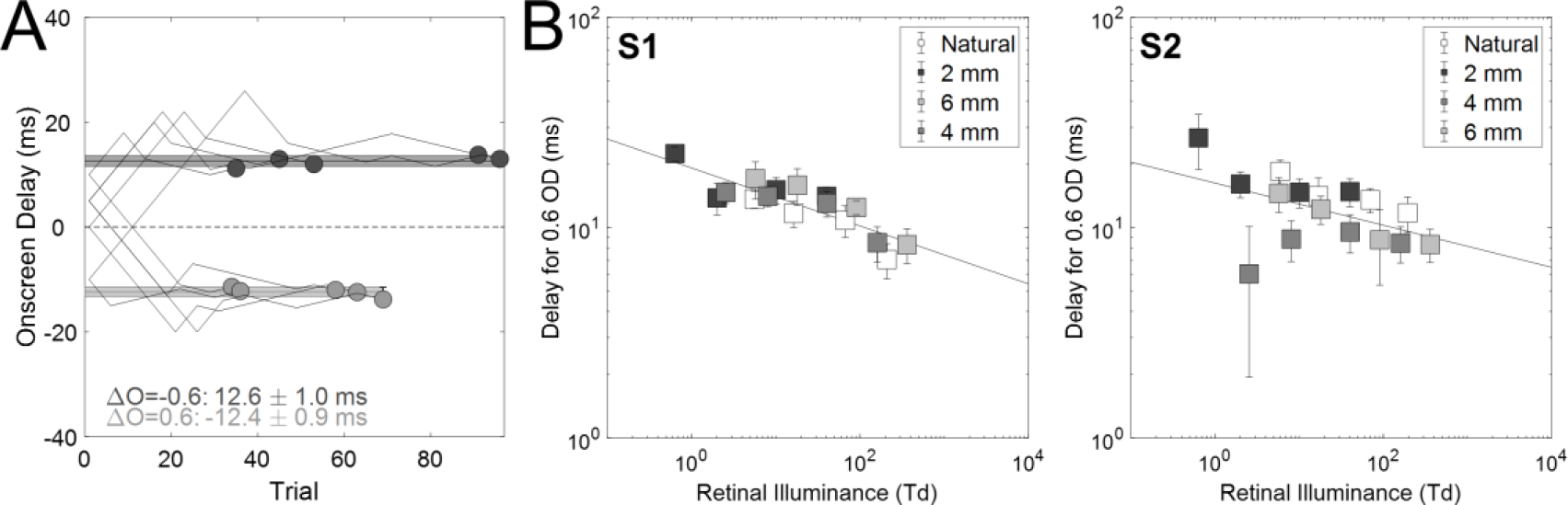
Classic Pulfrich effect results. **A**. Runs from the condition in which the left-eye image was 75% dimmer than the right-eye image are in dark gray (ΔO = -0.6 OD). Runs from the condition in which the right-eye image was 75% dimmer than the left-eye image are shown in light gray (ΔO = +0.6 OD). In both conditions, both eyes were sharply focused on the screen distance. Runs have different starting points, ranging from -15 ms to 15 ms. A shaded dark or light gray region represents the average and its width the standard deviation across runs, indicating the critical onscreen delay and the uncertainty for both conditions. **B**. Critical onscreen delay magnitude for every retinal illuminance level (abscissa) and pupil size (colors) for subject 1 (left subplot) and subject 2 (right subplot). Critical onscreen delay increases when the retinal illuminance decreases, and pupil size has no effect. The Classic Pulfrich effect is stronger when light-level is lower.

Figure 5B shows the impact of overall light-level on the strength of the Classic Pulfrich effect, with one panel for each observer. Clearly, larger interocular delays are associated with lower light levels, and smaller interocular delays are associated with higher light levels. For S1 (Figure 5B, left panel), the slope of the linear regression in the log-log space reflects the power of the power function, which is -0.14 ms/td (CI^68^ = [-0.16, -0.11]) for all pupil sizes. For S2 (Figure 5B, right panel) the slope is -0.10 ms/td (CI^68^ = [-0.15 -0.04]) for all pupil sizes. These results replicate findings reported by Lit (1949)^2^ and Prestrude (1971)^14^ (see Supplementary Figure S2).

Overall light-level has a similar impact on the strengths of the Reverse and Classic Pulfrich effect (see Figures 4 and 5). The most evident difference between the two experiments is that pupil size does not affect the Classic Pulfrich effect, whereas it has a systematic effect on the Reverse Pulfrich effect. These issues are discussed below.

## DISCUSSION

The Classic and the Reverse Pulfrich effects both increase in strength as the overall light-level decreases. That is, as overall light-level decreases, a fixed interocular difference in defocus blur or a fixed (proportional) interocular difference in retinal illuminance causes larger interocular differences in processing speed. The results for the Reverse Pulfrich effect are novel. The results reported for the Classic Pulfrich effect agree with previously published data.

A notable limitation of this study is its small sample size and the fact that both subjects were authors. The custom 4f tunable lens optical systems that we used were portable prototypes that necessitated an alignment procedure that was impractical to perform on naïve observers (see Methods). We note, however, that many articles on related aspects of visual processing, and that are now considered classics, report data from only one or two subjects that were often the authors of the study^2,16,17^. The facts that, in the present study, the data from one subject resembled the data from the other, that our measurements largely replicate those of Lit (1949) and Prestrude (1971)^14^, and that these classic papers have stood the test of time, all suggest the current results are likely to replicate. Nevertheless, to draw more robust conclusions, future research should be conducted on a larger number of subjects with more easily calibrated optical systems.

### The impact of overall light-level

Visual processing speed changes systematically with overall light-level: interocular delays decrease linearly on a log-log scale which entails that changes in processing speed are well-characterized by a power law. This finding holds true regardless of whether the interocular delays are induced by interocular differences in focus error (Figure 4B) or by interocular differences in luminance (Figure 5B). Changes in the temporal response properties of the retina almost certainly underlie these results.

Evidence indicates that the retina responds more sluggishly when overall light-level is lower. This evidence ranges from direct in vitro single-unit recordings^18^, to electroretinogram records^19^, to a range of tightly controlled psychophysical studies^14,17,20,21^. However, although the Classic Pulfrich effect has been attributed to retinal processes^14,15,20,22,23^, it seems unlikely that the Reverse Pulfrich effect can be attributed to the same physiological site. The Classic Pulfrich effect is caused by interocular differences in light-level. The Reverse Pulfrich effect is caused by different spatial-frequency content in the two eyes^3–5,24,25^. (The Reverse Pulfrich occurs when one eye’s image is more sharply focused than the other. The sharper eye is processed more slowly because it contains higher spatial frequencies^3^). Neural selectivity for different spatial frequencies does not emerge until early visual cortex^26^. As a consequence, spatial-frequency-based neural differences in processing speed most likely do not emerge until early visual cortex^27–29^. Hence, although the primary physiological sites at which the interocular neural delays first emerge that underlie the Classic and Reverse Pulfrich effects are likely to be retinal and cortical, respectively, the modulatory impact of overall light-level on the sizes of these effects can most likely be attributed to light-level-induced changes in the temporal properties of retinal response.

### The impact of pupil size

The custom 4f tunable lens system (see Figure 1D) provides precise control over pupil size because it projects a diaphragm of fixed size into the pupil plane of the observer. This aspect of our experimental design allowed us to isolate the modulatory impact of pupil size as overall light-level changed.

The manner in which pupil size modulates interocular delay is the most striking difference between the results in the two experiments. For the Classic Pulfrich effect, with well-focused images, the interocular delay associated with a given proportional difference in retinal illumination was unaffected by pupil size. This is to be expected. Although optical quality changes with pupil size, because both eyes were equally well focused (and because pupil sizes were the same in both eyes), changes in optical quality were matched between the eyes. Hence, at a given overall light-level, the only factor driving the interocular delays was the proportional differences in light level (|ΔO|=0.6 OD).

For the Reverse Pulfrich effect, the 4 mm, 6 mm, and natural pupil sizes—which ranged between 4 mm and 6 mm—resulted in larger interocular delays than the 2 mm pupil size. This result is expected from elementary geometric optics (see Figure 1 and Equation 5) and from previous findings in the literature regarding the relationship between retinal blur and processing speed^3,4^. The reasoning is as follows. Assuming geometric optics and no higher-order aberrations, the amount of retinal blur is linearly related to pupil size for a given focus error (Equation 5). Large pupils yield more defocus blur than small pupils for a given focus error (i.e., 3.0 D in these experiments). Bigger interocular differences in defocus blur cause larger interocular processing delays. So, a 3.0 D difference in focus error with large pupils is expected to generate larger interocular processing delays than a 3.0 D difference in focus error with small pupils.

The reasoning above accounts for why the 4 mm, 6 mm, and natural pupil sizes resulted in larger interocular delays than the 2 mm pupils. However, it does not account for why delays associated with the 4 mm and 6 mm pupils were so similar to one another (see Figure 4B). Other factors must be responsible. Defocus blur is not the only source of retinal blur in human eyes. Higher-order aberrations also contribute. Importantly, relative to defocus and astigmatism, the impact of higher-order aberrations on retinal blur increases with pupil size^30–33^. As a consequence, the presence of higher-order aberrations causes retinal blur to change less dramatically with pupil size than if they were absent. Hence, the similarity of the delays associated with the 4 mm, 6 mm, and natural pupil sizes may be due to higher-order aberrations. Development of a formal model that relates higher-order aberrations to blur discriminability—and hence, processing delay—may be a productive way forward and is left for future work.

### Monovision, overall light-level, and Anti-Pulfrich corrections

Previous research has examined how monovision-correction strengths impact the size of interocular processing delays and the severity of the resultant depth misperceptions caused by the Reverse Pulfrich effect^3–5^ can be large enough to cause safety concerns. These studies have found that interocular differences in processing speed are proportional to the interocular difference in dioptric power induced by monovision corrections over a wide range. In this study, we showed that decreasing overall light-level increases processing delays for a given interocular difference in focus error. These findings suggest that monovision corrections may pose an even more significant safety concern under low-light-level conditions (e.g., nighttime driving) than at high-light-level conditions.

In this study, we used focus differences of 3.0 D between the eyes, which creates optical conditions that are rarely induced by monovision prescriptions. We used this large difference in focus error because pilot data indicated that, of several tested focus-error differences, the 3.0 D difference produced the largest effect sizes (see footnote 1).

Anti-Pulfrich monovision corrections are aimed at eliminating the depth misperceptions, and hence the safety concerns, caused by the interocular differences in defocus blur induced by regular monovision corrections^3^. Anti-Pulfrich monovision corrections take advantage of the fact that the Classic and Reverse Pulfrich effects have opposite signs. That is, darkening an image slows down how fast it is processed; blurring an image speeds up how fast it is processed. At a particular overall light-level, the Reverse Pulfrich effect that is caused by a given blur difference can thus be eliminated by appropriately tinting the lens of one eye^3,4^.

In the current study, we showed that both the Reverse and the Classic Pulfrich effects increase in strength with decreases in overall light-level. However, these changes in effect size occur at different rates (see slopes in Figures 4B and 5B). More specifically, the Classic Pulfrich effect changes approximately twice as fast as the Reverse Pulfrich effect with overall light-level. Hence, a difference in tint that is effective at one light-level would not be effective at another light-level. Thus, to develop an effective anti-Pulfrich monovision correction for all light levels, the tint difference would need to change with light-level. Photochromic contact lenses^34,35^ change their transmittance with ambient light-level. This technology may enable an appropriate delivery system for an anti-Pulfrich monovision correction that is effective at all overall light-levels.

## CONCLUSION

In this study, we report that the severity of the Reverse Pulfrich effect increases when overall light-level decreases: the same interocular focus difference causes larger depth misperceptions in nightlight than in daylight. We also replicate multiple studies showing that the severity of the Classic Pulfrich is similarly affected. These results motivate a full characterization of how light level interacts with other optical factors (e.g., higher-order aberrations) likely to impact the Reverse (and Classic) Pulfrich effect(s). Optical technologies like anti-Pulfrich monovision corrections that aim to eliminate the depth misperceptions caused by typical monovision corrections, must account for how the Reverse and Classic Pulfrich effects change with light-level.

## Acknowledgments

We thank Jessica Morgan and Yu You Jiang at Scheie Eye Institute, University of Pennsylvania for technical assistance. This research was supported by “la Caixa” Foundation (ID 100010434; LCF/BQ/DR19/11740032) to V.R.L., and NIH grant R01-EY028571 from the National Eye Institute & Office of Behavioral and Social Science Research to J.B. The custom 4f optical system was developed at the Institute of Optics in Madrid, Spain. The psychophysical experiments were conducted at the University of Pennsylvania in Philadelphia, PA.

## Measurements of pupil size

Supplementary Figure 1 shows the pupil size measured for each overall light-level.

### Comparison to classic literature

**Figure S1.**
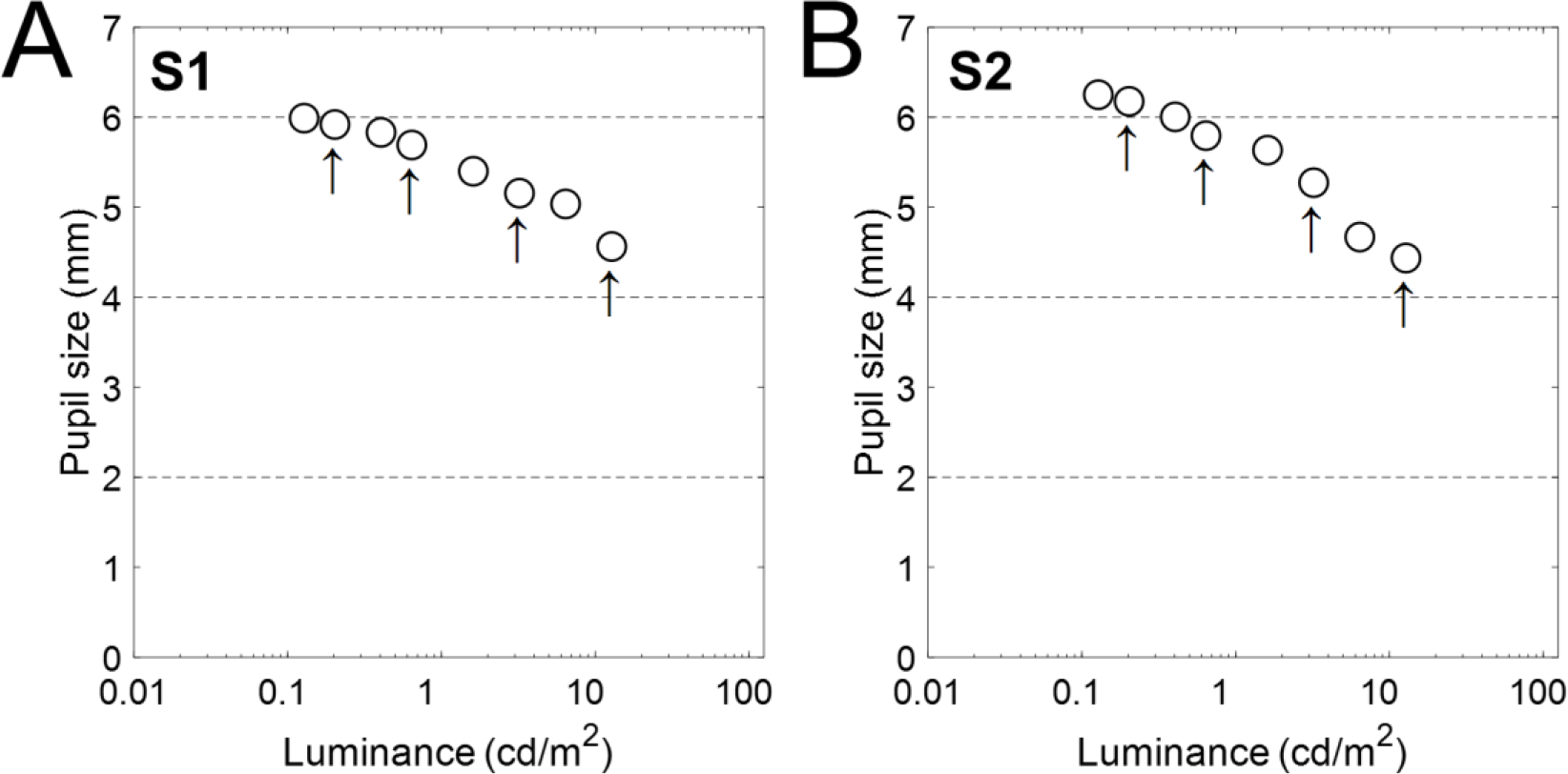
Pupil measurements. Pupil size measured as a function of overall light-level (luminance level emitted by the display) for natural pupil condition. Arrows mark the overall light-level conditions measured in the experiments. Horizontal dashed lines indicate the pupil diameters that were fixed pupil conditions in the experiments (i.e., 2, 4, and 6 mm). **A**. Subject S1. **B**. Subject S2.

For the Classic Pulfrich effect, there were data available from other experiments that already performed measurements on how the Classic Pulfrich changes with overall light-level. In both Lit^2^ and Prestrude^14^, they measured the neural delay for interocular differences in light levels up to ΔO=3.0 OD and for retinal illuminance levels ranging from 2.3 to 2500 trolands (in our measurements we measure ΔO=±0.6 OD and retinal illuminance levels from 0.6 to 360 trolands). Lit measured the perceived depth in distance, and later transformed into a neural temporal delay in two subjects. Prestrude directly measured the neural delay in four subjects. For each luminance level, they measured several interocular luminance differences. However, for interocular differences higher than ΔO = 1.0 OD, the neural delay began to behave non-linearly. Besides, we measured luminance differences of ΔO=0.6 OD. For a fair comparison with the methodology of our study, we calculated the neural delay caused by a ΔO=0.6 OD filter for each retinal illuminance level. Then, we estimated the linear regression of the neural delays measured up to ΔO=1.0 OD in their studies. Supplementary Figure S2A and S2B shows these estimations. Supplementary Figure S2C shows the delay as a function of retinal illuminance for both studies from the literature. The slope of the linear regression in the log-log space reflects the power of the power function (see Equation 5), which is -0.23 ms/td for Lit 1949 and -0.24 ms/td for Prestrude 1971.

**Figure S2.**
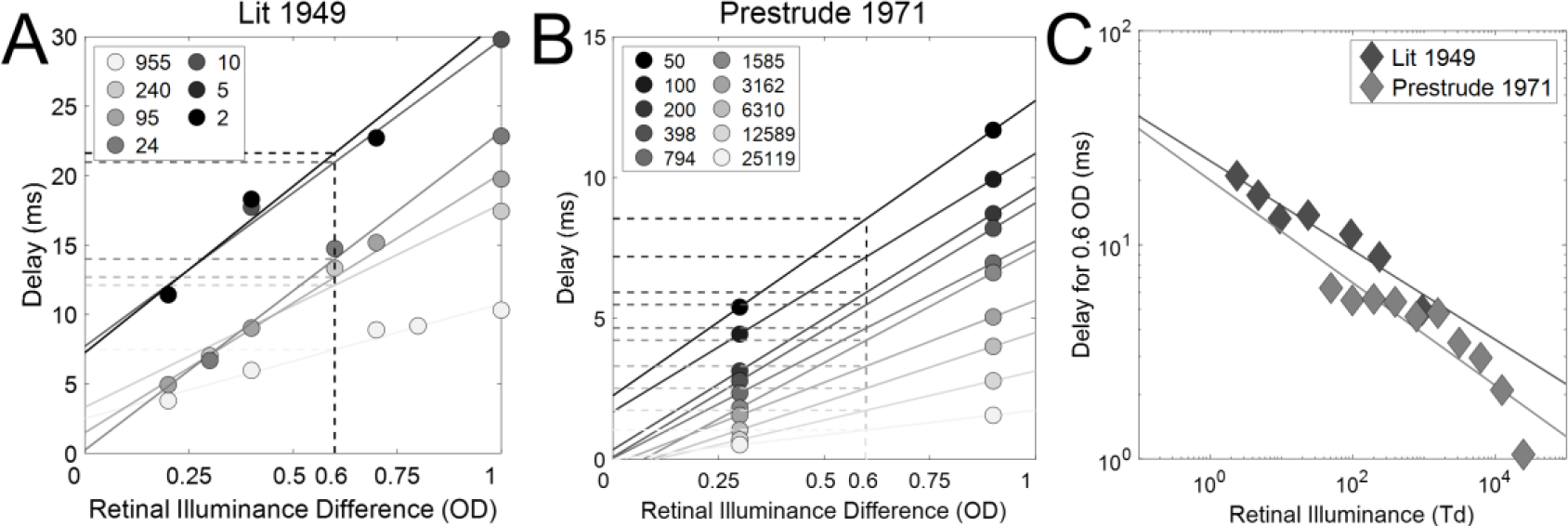
Classic Pulfrich effect for different overall light-levels from the literature. **A**. Estimation of the delay for ΔO=0.6 OD from Lit 1949. **B**. Estimation of the delay for ΔO=0.6 OD from Prestrude 1971. **C**. Delay for a filter of 0.6 OD in ms for every retinal illuminance level measured in Lit 1949 and Prestrude 1971, averaged across subjects.

## Footnotes

1 We noted above that monovision corrections typically induce focus-error differences of 1.5 D, and rarely exceed differences of 2.5 D. One might have a concern, therefore, that measurements reported here are not reflective of typical monovision prescriptions. We used 3.0D differences because, in pilot experiments, they produced the largest effect sizes. Pilot data also suggested that interocular delay scales linearly with focus error up to 3.0 D. This informal observation is consistent with published observations that the Reverse Pulfrich effect scales linearly with differences in focus error up to 1.5 D (no data on larger focus-error differences has been published). If this latter observation holds, the interocular delay associated with a 3.0 D focus error could simply be divided by two to obtain the interocular delay associated with 1.5D, the typical monovision correction strength (see Discussion).

