## Supplementary Figure for "The impact of overall light-level on the reverse Pulfrich effect"

### Running title: Overall light-level and the reverse Pulfrich effect

Victor Rodriguez-Lopez, MSc[1,2,*],
Benjamin Chin, PhD[2],
Johannes Burge, PhD[2,3,4]

[1.] Institute of Optics, Spanish National Research Council (IO-CSIC), Madrid, Spain
[2.] Department of Psychology, University of Pennsylvania, Pennsylvania PA
[3.] Neuroscience Graduate Group, University of Pennsylvania, Pennsylvania PA
[4.] Bioengineering Graduate Group, University of Pennsylvania, Pennsylvania PA

*Measurements of pupil size*

Supplementary Figure 1 shows the pupil size measured for each overall light-level.

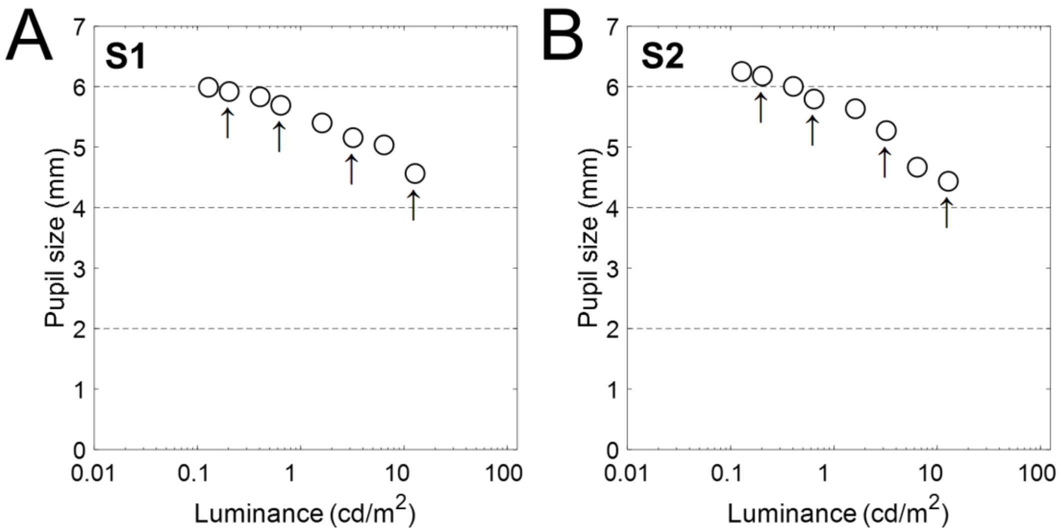


**Figure S1. Pupil measurements**. Pupil size measured as a function of overall light-level (luminance level emitted by the display) for natural pupil condition. Arrows mark the overall light-level conditions measured in the experiments. Horizontal dashed lines indicate the pupil diameters that were fixed pupil conditions in the experiments (i.e., 2, 4, and 6 mm). **A.** Subject S1. **B.** Subject S2.

*Comparison to classic literature*

For the Classic Pulfrich effect, there were data available from other experiments that already performed measurements on how the Classic Pulfrich changes with overall light-level. In both Lit[2] and Prestrude[14], they measured the neural delay for interocular differences in light levels up to ΔO = 3.0 OD and for retinal illuminance levels ranging from 2.3 to 2,500 trolands (in our measurements we measure ΔO = ±0.6 OD and retinal illuminance levels from 0.6 to 360 trolands). Lit measured the perceived depth in distance and later transformed into a neural temporal delay in two subjects. Prestrude directly measured the neural delay in four subjects. For each luminance level, they measured several interocular luminance differences. However, for interocular differences higher than ΔO = 1.0 OD, the neural delay began to behave non-linearly. Besides, we measured luminance differences of ΔO=0.6 OD. For a fair comparison with the methodology of our study, we calculated the neural delay caused by a ΔO = 0.6 OD filter for each retinal illuminance level. Then, we estimated the linear regression of the neural delays measured up to ΔO=1.0 OD in their studies. Supplementary Figure S2A and S2B shows these estimations. Supplementary Figure S2C shows the delay as a function of retinal illuminance for both studies from the literature. The slope of the linear regression in the log-log space reflects the power of the power function (see Equation 5), which is -0.23 ms/td for Lit 1949 and -0.24 ms/td for Prestrude 1971.

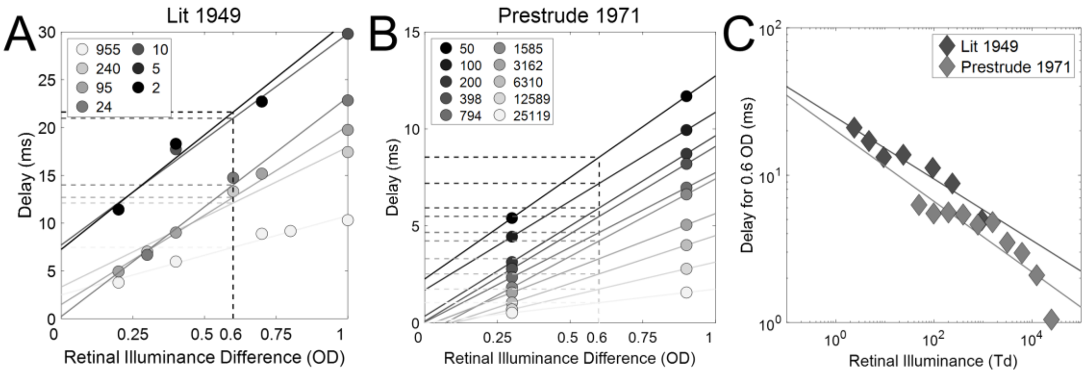


**Figure S2. Classic Pulfrich effect for different overall light-levels from the literature. A.** Estimation of the delay for ΔO=0.6 OD from Lit 1949. **B.** Estimation of the delay for ΔO=0.6 OD from Prestrude 1971. **C.** Delay for a filter of 0.6 OD in ms for every retinal illuminance level measured in Lit 1949 and Prestrude 1971, averaged across subjects.
